# Identifying Pleiotropic Effects: A Two-Stage Approach Using Genome-Wide Association Meta-Analysis Data

**DOI:** 10.1101/184895

**Authors:** Xing Chen, Yi-Hsiang Hsu

**Affiliations:** Molecular and Integrative Physiological Sciences, Harvard School of Public Health, Boston, MA; Hebrew SeniorLife Institute for Aging Research and Harvard Medical School, Boston, MA; BROAD Institute of MIT and Harvard, Boston, MA

## Abstract

Pleiotropic effects occur when a single genetic variant independently influences multiple phenotypes. In genetic epidemiological studies, multiple endo-phenotypes or correlated traits are commonly tested separately in a univariate statistical framework to identify associations with genetic determinants. Subsequently, a simple look-up of overlapping univariate results is applied to identify pleiotropic genetic effects. However, this strategy offers limited power to detect pleiotropy. In contrast, combining correlated traits into a composite test provides a powerful approach for detecting pleiotropic genes. Here, we propose a two-stage approach to identify potential pleiotropic effects by utilizing aggregated results from large-scale genome-wide association (GWAS) meta-analyses. In the first stage, we developed two novel approaches (direct linear combining, dLC; and empirical combining, eLC) combining correlated univariate test statistics to screen potential pleiotropic variants on a genome-wide scale, using either individual-level or aggregated data. Our simulations indicated that dLC and eLC outperform other popular multivariate approaches (such as principal component analysis (PCA), multivariate analysis of variance (MANOVA), canonical correlation (CCA), generalized estimation equations (GEE), linear mixed effects models (LME) and O’Brien combining approach). In particular, eLC provides a notable increase in power when the genetic variant exhibits both protective and deleterious effects. In the second stage, we developed a unique approach, conditional pleiotropy testing (cPLT), to examine pleiotropic effects using individual-level data for candidate variants identified in Stage 1. Simulation demonstrated reduced type 1 error for cPLT in identifying pleiotropic genetic variants compared to the typical conditional strategy. We validated our two-stage approach by performing a bivariate GWA study on two correlated quantitative traits, high-density lipoprotein (HDL) and triglycerides (TG), in the Genetic Analysis Workshop 16 (GAW16) simulation dataset. In summary, the proposed two-stage approach allows us to leverage aggregated summary statistics from univariate GWAS and improves the power to identify potential pleiotropy while maintaining valid false-positive rates.

**Author Summary:** Pleiotropy, occurring when a single genetic variant contributes to multiple phenotypes, remains difficult to identify in genome-wide association studies (GWAS). To leverage data for multiple phenotypes and incorporate univariate GWAS summary results, we propose a novel two-stage approach for discovering potential pleiotropic variants. In the first stage, two novel combining approaches were developed to screen potential pleiotropic variants on a genome-wide scale. Simulations demonstrated the superior statistical power of these approaches over other multivariate methods. In the second stage, our approach was used to identify potential pleiotropy in the candidate marker sets generated from the first stage. The proposed two-stage approach was applied to the GAW16 simulation dataset to discover pleiotropic variants associated with high-density lipoprotein and triglycerides. In summary, we demonstrate that the proposed two-stage approach can be applied as a viable and robust strategy to accommodate phenotypic and genetic heterogeneity for discovering potential pleiotropy on genome-wide scale.

## Introduction

Genome-wide association studies (GWAS) have been successfully identified genetic determinants underlying complex human diseases [1]. Conventionally, univariate analytic approaches were used to study associations between numerous genetic variants and a single phenotype, one at a time, although multiple correlated phenotypes were often available for joint examination [2–5]. Pleiotropic effects, in which genetic variants influence more than one phenotype or disease, have been widely observed in recent GWAS findings [6]. For example, a genetic variant in the *glucokinase regulator* gene (*GKCR*) is associated with increased concentrations of plasma triglycerides but reduced fasting glucose levels [7]. Other notable findings include two single-nucleotide polymorphisms (SNPs) in intron 1 of *SRY-box 6* (*SOX6*) that are associated with both body-mass index (BMI) and hip bone mineral density (BMD) in adult Caucasians [8], and a region on chromosome 8q24 associated with prostate, breast, and colorectal cancers and Crohn’s disease in different populations [9–12].

Multivariate methods by analyzing correlated traits jointly provide statistical advantages in increasing power and accuracy of parameter estimation in linkage studies [13–16]. When individual-level data are available, a few multivariate methods have been developed to jointly analyze correlated traits on a genome-wide scale [14,17–21]. These classic multivariate methods include principle component analysis (PCA) [22], multivariate analysis of variance (MANOVA), generalized estimation equations (GEE) [23–26], and linear mixed effect models (LME) [24]. Additionally, extended GEE has been incorporated in family-based tests (FBAT) [20] for analyzing correlated subjects. With the increasing abundance of GWAS data, meta-analyses have become popular for pooling results from multiple cohorts to increase the sample size and power of identifying genetic determinants that underlie the etiology of complex disease [27]. However, most multivariate methods described above require access to individual-level data, limiting their use in large-scale GWAS meta-analyses with only available summary statistics [28–30] and suggesting alternative statistical approaches are needed to improve the detection of potential pleiotropic effects.

In contrast to the classical multivariate methods, approaches to directly combine test statistics by integrating summary statistics from multiple univariate meta-analyses into a global test, may overcome the limitations of classical multivariate methods. Fisher first proposed the combined probability test, which combines results from several independent tests of the same overall hypothesis (H_0_) [31]. However, Fisher’s exact test method likely introduces inflated type-1 error rates when tests are dependent [32]. O’Brien and Wei separately proposed better approaches [33–36] to combine test statistics from correlated traits. Although demonstrating greater power when individual test statistics are homogeneous, their methods usually do not achieve desired power when the effect directions are different [37]. Yang et al. [37] extended O’Brien’s approach by using sample splitting and cross-validation methods to gain power when heterogeneous genetic variants exist. However, individual-level data are required to estimate the optimal weight in this approach and only a subset of study samples is used in inferring the final test statistics, limiting the applications of Yang’s method. Another approach, named “TATES”, was recently proposed to combine p-values from univariate GWAS while correcting the observed phenotypic correlations [38]. This approach was not explicitly examined for traits that are negatively correlated. Province et al. [39] also provide a powerful tool to conduct multivariate analyses on correlated phenotypes with estimated weighting by tetrachoric correlations; however, it remains unclear whether their method achieves better power than others, especially when mixed genetic effect directions exists.

Notably, significant findings from multivariate analyses do not always indicate pleiotropic effects. Such findings could be likely introduced by correlations between two correlated phenotypes, rather than by independent effects of a genetic variant [40]. Several methods have been proposed to differentiate between causal genetic effects and the secondary associations caused by correlations between phenotypes [40,41]. However, few methods have been studied and applied in the context of pleiotropy testing.

To overcome these limitations, we propose a two-stage approach (Figure 1). In the first-stage (genome-wide screening), two methods, direct linear combining approach (dLC) and empirical linear combining approach (eLC), using aggregated results from meta-analysis were developed. In the second-stage, pleiotropy was identified in markers selected from the first stage using the conditional pleiotropy testing (cPLT) approach. We first described statistical power and type 1 error our proposed methods comparing to the power and type 1 error of our existing methods by simulation. We then further demonstrated the proposed two-stage approach to identify pleiotropy in the Genetic Analysis Workshop 16 (GAW16) Framingham heart Study data sets.

**Figure 1.**
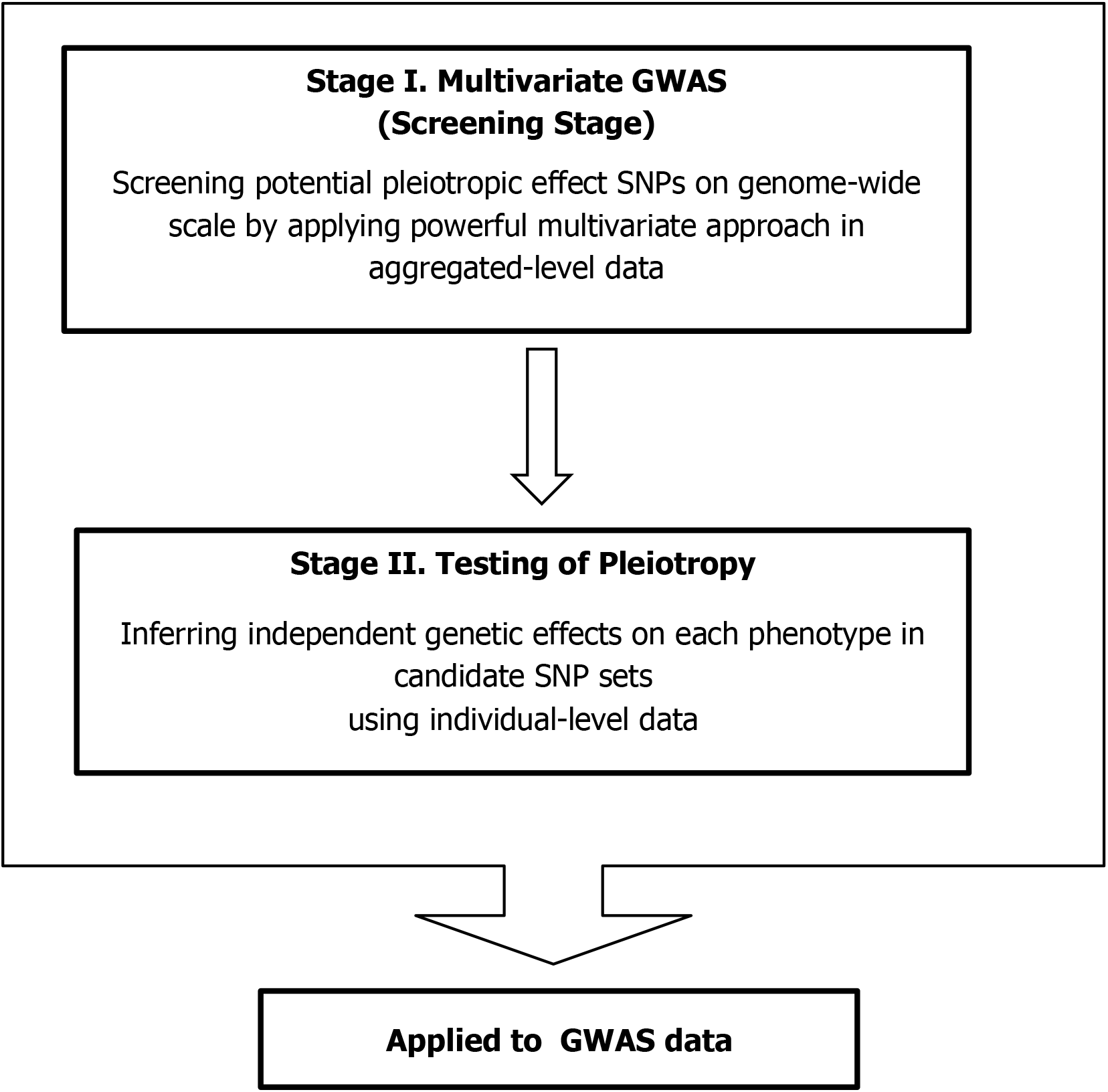
Study design of proposed two-stage approach. Simulation was conducted to estimate the performance of proposed multivariate GWAS approaches and testing of pleiotropy method. The proposed approaches were then applied in a GWAS dataset with correlated serum lipid measurements (HDL and TG).

## Results

We conducted extensive simulations to evaluate the performance of our proposed two-stage strategy, and compared the results with existing methods. Key simulation parameters are provided in Table 1.

**Table 1.** Key parameters of simulation scenarios

| Series | Note | Scenario | $H_1$ | Phenotype 1 ($Y_1$) | | Phenotype 2 ($Y_2$) | |
| --- | --- | --- | --- | --- | --- | --- | --- |
| | | | | $H_2$ | $\beta$ | $H_2$ | $\beta$ |
| I. Higher effect size | Pleiotropy with equal effect | A | $\beta_1 = \vert \beta_2\vert > 0$ | 0.02 | 0.34 | 0.02 | 0.34 |
| | Pleiotropy with unequal effect | B | $\beta_1 > \vert \beta_2\vert > 0$ | 0.02 | 0.34 | 0.01 | 0.24 |
| | Non-pleiotropy | C | $\beta_1 > 0, \beta_2 = \beta_1 + \varepsilon$ | 0.02 | 0.34 | random noise | |
| | Pleiotropy on multiple phenotypes | D<sup>a</sup> | $\beta_1 = \vert \beta_2\vert = \vert \beta_3\vert > 0$ | 0.02 | 0.34 | | |
| II. Lower effect size | Pleiotropy with equal effect | A | $\beta_1 = \vert \beta_2\vert > 0$ | 0.005 | 0.17 | 0.005 | 0.17 |
| | Pleiotropy with unequal effect | B | $\beta_1 > \vert \beta_2\vert > 0$ | 0.005 | 0.17 | 0.001 | 0.075 |
| | Non-pleiotropy | C | $\beta_1 > 0, \beta_2 = \beta_1 + \varepsilon$ | 0.005 | 0.17 | random noise | |
| | Pleiotropy on multiple phenotypes | D<sup>b</sup> | $\beta_1 = \vert \beta_2\vert = \vert \beta_3\vert > 0$ | 0.005 | 0.17 | | |

Note: Minor Allele Frequency=10% for all scenarios.

Note: Minor Allele Frequency=10% for all scenarios.

Variance of environmental effects is fixed at 1 for all scenarios.

Total of 1,000 subjects were simulated in each replicate.

^a^ Equal heritability in this scenario is assigned as 0.02 for all three quantitative traits.

^b^ Equal heritability in this scenario is assigned as 0.005 for all three quantitative traits.

### First-Stage (GWAS Screening): Performance of multivariate methods

Using **individual-level of genotype and phenotype data**, valid type I error rates, as expected, were consistently observed across all simulation scenarios at different nominal levels for all examined multivariate methods (Table 2). In particular, our proposed methods, dLC and eLC, demonstrate robustness under the null hypothesis, regardless of the directions of phenotypic residual correlation and estimations of the covariance matrix **Σ**. The estimated power is presented in Table 3. Simulation Series I assessed the performance of various multivariate approaches under relevant hypotheses, in which relatively large genetic effects were simulated. The results are presented with respected to two alternative hypotheses, H_1_:β_1_=|β|_2_>0 and H_1_:β_1_>|β_2_|>0. In particular, we assigned a mixture of protective and deleterious genetic effects when negative phenotypic residual ρ was adopted in the simulation. **All the methods performed well when test statistics were homogeneous, but their power varied considerably under the hypothesis of mixed genetic effects and effect directions**. For instance, our proposed approaches showed comparable power to MANOVA under the β1=|β|2>0 alternative hypothesis; GEE, LME, and the O’Brien method (OB) had lower power. When traits were highly correlated, principle component analysis (PCA) outperformed all other methods; however, when traits were barely related (ρ=0.01), the PCA showed less power than our proposed dLC and eLC methods. The results under the β1>|β2|>0 alternative hypothesis produced similar conclusions across various simulation scenarios. In general, our proposed mthods, dLC and eLC, are substantially superior to univariate analyses (look-up the overlapping among multiple univariate GWAS signals) and other classical multivariate methods under all alternative hypotheses. Additionally, eLC outperformed dLC marginally in this simulation series.

**Table 2.**
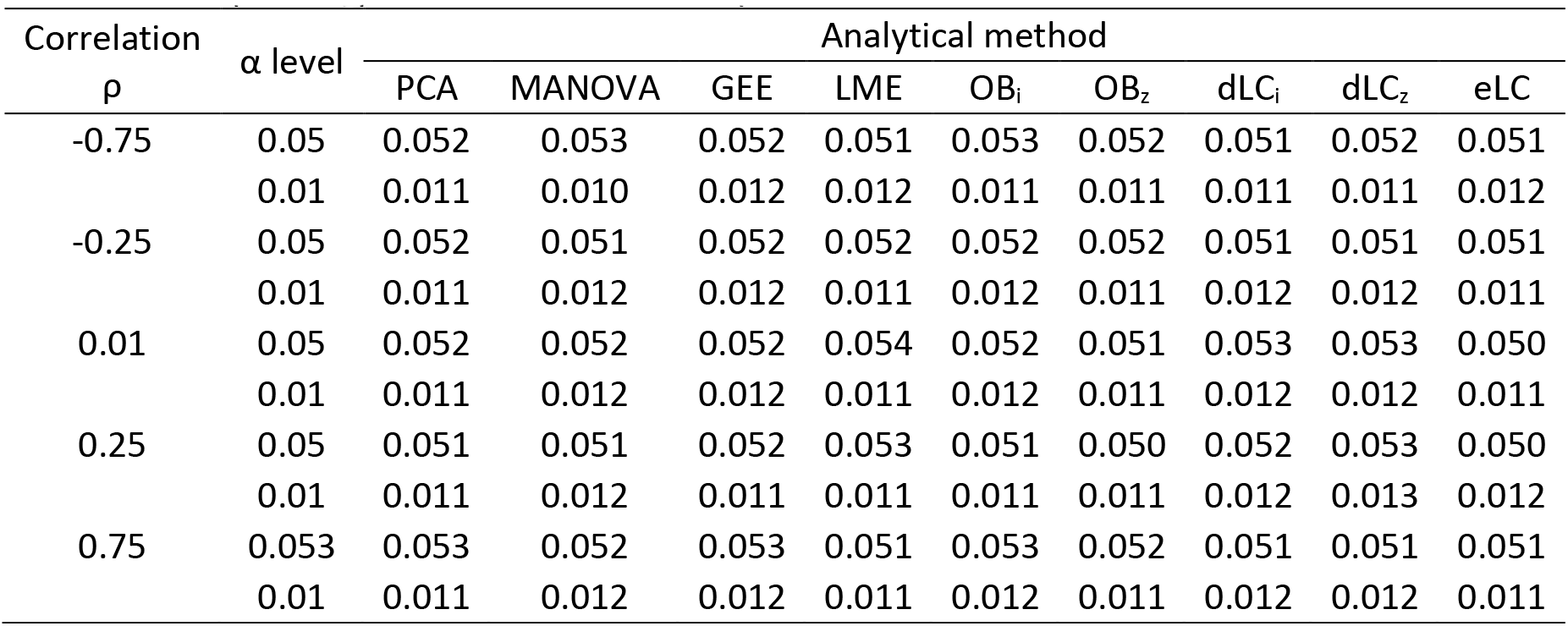
Estimated type I error rates of multivariate approaches in 1000 replicates with different phenotypic residual correlations ρ

| Correlation<br>$\rho$ | $\alpha$ level | Analytical method | | | | | | | | |
| --- | --- | --- | --- | --- | --- | --- | --- | --- | --- | --- |
|  |  | PCA | MANOVA | GEE | LME | OB<sub>i</sub> | OB<sub>z</sub> | dLC<sub>i</sub> | dLC<sub>z</sub> | eLC |
| -0.75 | 0.05 | 0.052 | 0.053 | 0.052 | 0.051 | 0.053 | 0.052 | 0.051 | 0.052 | 0.051 |
|  | 0.01 | 0.011 | 0.010 | 0.012 | 0.012 | 0.011 | 0.011 | 0.011 | 0.011 | 0.012 |
| -0.25 | 0.05 | 0.052 | 0.051 | 0.052 | 0.052 | 0.052 | 0.052 | 0.051 | 0.051 | 0.051 |
|  | 0.01 | 0.011 | 0.012 | 0.012 | 0.011 | 0.012 | 0.011 | 0.012 | 0.012 | 0.011 |
| 0.01 | 0.05 | 0.052 | 0.052 | 0.052 | 0.054 | 0.052 | 0.051 | 0.053 | 0.053 | 0.050 |
|  | 0.01 | 0.011 | 0.012 | 0.012 | 0.011 | 0.012 | 0.011 | 0.012 | 0.012 | 0.011 |
| 0.25 | 0.05 | 0.051 | 0.051 | 0.052 | 0.053 | 0.051 | 0.050 | 0.052 | 0.053 | 0.050 |
|  | 0.01 | 0.011 | 0.012 | 0.011 | 0.011 | 0.011 | 0.011 | 0.012 | 0.013 | 0.012 |
| 0.75 | 0.053 | 0.053 | 0.052 | 0.053 | 0.051 | 0.053 | 0.052 | 0.051 | 0.051 | 0.051 |
|  | 0.01 | 0.011 | 0.012 | 0.012 | 0.011 | 0.012 | 0.011 | 0.012 | 0.012 | 0.011 |

Note: Sample Size N=1000.

Note: Sample Size N=1000.

Subscript *i* indicates that individual-level data were used; *z* indicates that aggregated data were used.

**Table 3.**
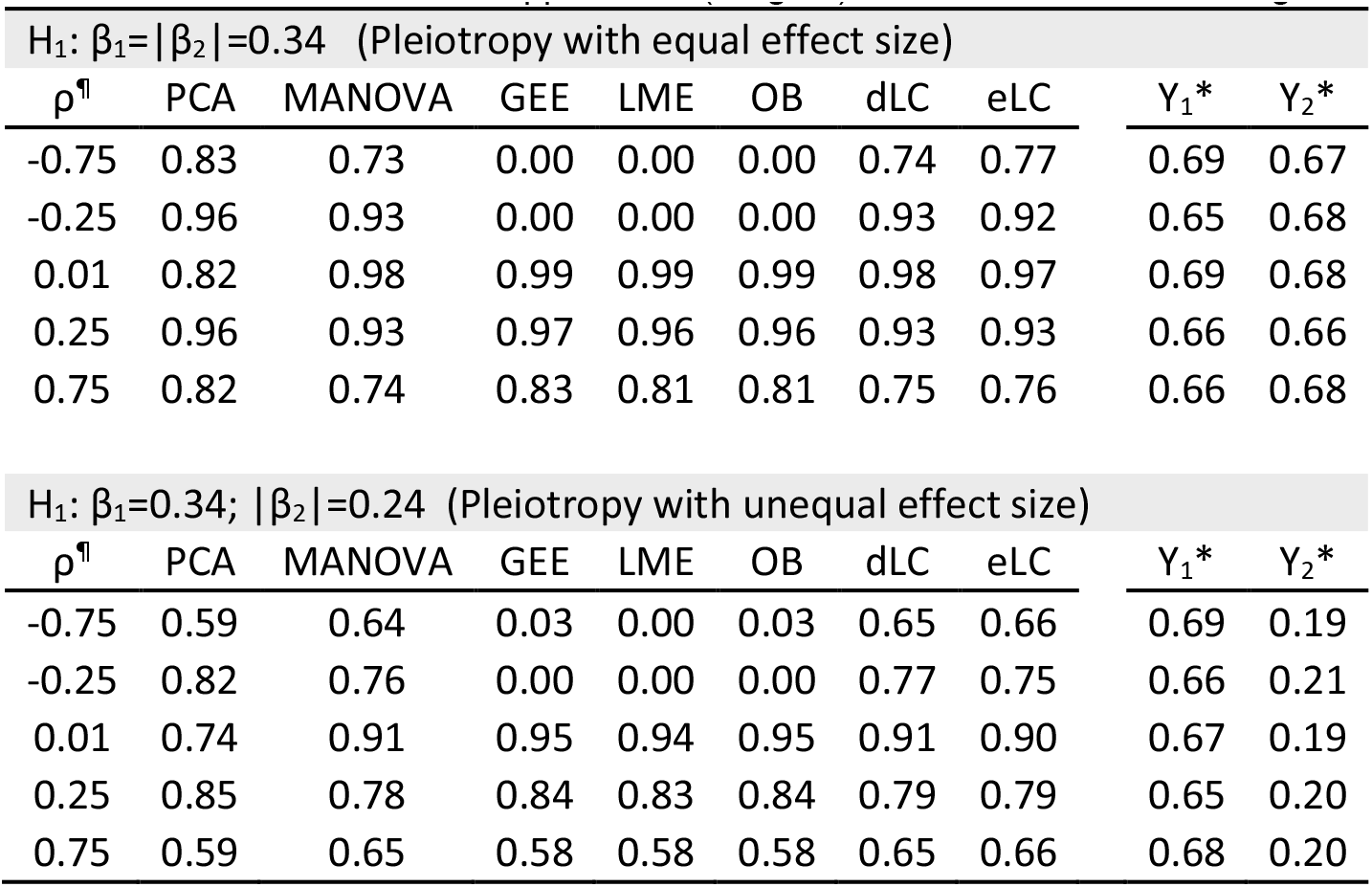
Power of multivariate approaches (Stage 1) in Simulation Series I using individual-level data.

| H<sub>1</sub>: $\beta_1 = \vert \beta_2\vert = 0.34$ (Pleiotropy with equal effect size) | | | | | | | | | |
| --- | --- | --- | --- | --- | --- | --- | --- | --- | --- |
| $\rho^{\text{¶}}$ | PCA | MANOVA | GEE | LME | OB | dLC | eLC | Y<sub>1</sub>* | Y<sub>2</sub>* |
| -0.75 | 0.83 | 0.73 | 0.00 | 0.00 | 0.00 | 0.74 | 0.77 | 0.69 | 0.67 |
| -0.25 | 0.96 | 0.93 | 0.00 | 0.00 | 0.00 | 0.93 | 0.92 | 0.65 | 0.68 |
| 0.01 | 0.82 | 0.98 | 0.99 | 0.99 | 0.99 | 0.98 | 0.97 | 0.69 | 0.68 |
| 0.25 | 0.96 | 0.93 | 0.97 | 0.96 | 0.96 | 0.93 | 0.93 | 0.66 | 0.66 |
| 0.75 | 0.82 | 0.74 | 0.83 | 0.81 | 0.81 | 0.75 | 0.76 | 0.66 | 0.68 |

| H<sub>1</sub>: $\beta_1 = 0.34$; $\vert \beta_2\vert = 0.24$ (Pleiotropy with unequal effect size) | | | | | | | | | |
| --- | --- | --- | --- | --- | --- | --- | --- | --- | --- |
| $\rho^{\text{¶}}$ | PCA | MANOVA | GEE | LME | OB | dLC | eLC | Y<sub>1</sub>* | Y<sub>2</sub>* |
| -0.75 | 0.59 | 0.64 | 0.03 | 0.00 | 0.03 | 0.65 | 0.66 | 0.69 | 0.19 |
| -0.25 | 0.82 | 0.76 | 0.00 | 0.00 | 0.00 | 0.77 | 0.75 | 0.66 | 0.21 |
| 0.01 | 0.74 | 0.91 | 0.95 | 0.94 | 0.95 | 0.91 | 0.90 | 0.67 | 0.19 |
| 0.25 | 0.85 | 0.78 | 0.84 | 0.83 | 0.84 | 0.79 | 0.79 | 0.65 | 0.20 |
| 0.75 | 0.59 | 0.65 | 0.58 | 0.58 | 0.58 | 0.65 | 0.66 | 0.68 | 0.20 |

¶: opposite effect direction was assigned when negative phenotypic correlation $\rho$ was used.

*: p-values were adjusted using Bonferroni corrections in these univariate analyses.

^¶^: opposite effect direction was assigned when negative phenotypic correlation ρ was used.

**For aggregated summary statistics**, we examined to what extent power is changed by applying our proposed methods under the same alternative hypotheses. MANOVA, PCA, GEE, and LME are unable to applied when only aggregative summary results are available. Thus, we only compared power between OB, dLC, and eLC (Table 4). OB was inferior when the univariate summary test statistics were heterogeneous. dLC and eLC were not only superior to OB when directions of effects were opposite, but nearly as powerful as OB when genetic effects are homogenous. Moreover, similar power was consistently observed by applying two distinct methods to estimate covariance matrix **Σ**. Again, eLC and dLC also showed significant advantages over univariate methods in detecting pleiotropic effects. Similar conclusions were drawn on the results from Simulation Series II, where smaller genetic effects were simulated (see **Supplementary Tables S1** and **S2**). These results demonstrate that our proposed methods have comparable power to other approaches when all effects are in the same direction and much greater power when genetic effects and directions are mixed. Our proposed methods can also applied in aggregated data (such as results from GWAS metaanalyses), thus efficiently increasing the sample size and power.

**Table 4.** Power of multivariate approaches (Stage 1) in Simulation Series I using aggregated data

| Alternative Hypothesis ($H_1$) | $\rho^{\P}$ | OB | dLC | eLC | $Y_1$ | $Y_2$ | $Y_1^*$ | $Y_2^*$ |
| --- | --- | --- | --- | --- | --- | --- | --- | --- |
| $\beta_1 = \vert \beta_2\vert = 0.34$ | -0.75 | 0.00 | 0.74 | 0.78 | 0.74 | 0.73 | 0.69 | 0.67 |
|  | -0.25 | 0.00 | 0.93 | 0.91 | 0.71 | 0.74 | 0.65 | 0.68 |
|  | 0.01 | 0.99 | 0.98 | 0.97 | 0.76 | 0.73 | 0.69 | 0.68 |
|  | 0.25 | 0.97 | 0.93 | 0.92 | 0.73 | 0.72 | 0.66 | 0.66 |
|  | 0.75 | 0.82 | 0.75 | 0.76 | 0.72 | 0.74 | 0.66 | 0.68 |
| $\beta_1 = 0.34; \vert \beta_2\vert = 0.24$ | -0.75 | 0.02 | 0.65 | 0.63 | 0.74 | 0.23 | 0.69 | 0.19 |
|  | -0.25 | 0.00 | 0.75 | 0.72 | 0.71 | 0.25 | 0.65 | 0.21 |
|  | 0.01 | 0.97 | 0.94 | 0.95 | 0.73 | 0.24 | 0.69 | 0.19 |
|  | 0.25 | 0.85 | 0.79 | 0.76 | 0.71 | 0.25 | 0.66 | 0.20 |
|  | 0.75 | 0.60 | 0.65 | 0.63 | 0.73 | 0.24 | 0.66 | 0.20 |
| $\beta_1 = \vert \beta_2\vert = 0.17$ | -0.50 | 0.02 | 0.44 | 0.43 | 0.39 | 0.39 | 0.31 | 0.31 |
|  | -0.25 | 0.01 | 0.52 | 0.50 | 0.38 | 0.39 | 0.28 | 0.32 |
|  | 0.01 | 0.74 | 0.62 | 0.63 | 0.37 | 0.35 | 0.28 | 0.27 |
|  | 0.25 | 0.58 | 0.46 | 0.45 | 0.34 | 0.34 | 0.26 | 0.27 |
|  | 0.50 | 0.50 | 0.40 | 0.41 | 0.36 | 0.37 | 0.27 | 0.30 |
| $\beta_1 = 0.17; \vert \beta_2\vert = 0.075$ | -0.50 | 0.11 | 0.30 | 0.27 | 0.37 | 0.07 | 0.29 | 0.04 |
|  | -0.25 | 0.07 | 0.29 | 0.31 | 0.35 | 0.06 | 0.27 | 0.04 |
|  | 0.01 | 0.39 | 0.35 | 0.35 | 0.37 | 0.07 | 0.28 | 0.04 |
|  | 0.25 | 0.29 | 0.29 | 0.30 | 0.35 | 0.06 | 0.29 | 0.04 |
|  | 0.50 | 0.23 | 0.29 | 0.29 | 0.35 | 0.05 | 0.28 | 0.04 |

<sup>¶</sup>: opposite effect direction was assigned when negative phenotypic correlation $\rho$ was used.

*: p-values were adjusted using Bonferroni correction in these univariate analyses.

^¶^: opposite effect direction was assigned when negative phenotypic correlation ρ was used.

**Multivariate analyses with more than 2 phenotypes**: We extended our proposed approaches from bivariate analysis (2 phenotypes) to multivariate analysis combining three correlated traits/phenotypes together. We generated data in the same manner as for the bivariate analysis under the null hypothesis, H_0_: β_1_=β_2_=β_3_=0, and two alternative hypotheses separately, H_1_: β_1_=-β_2_=-β_3_ >0 and H_1_: β_1_=β_2_=-β_3_ >0. We excluded PCA in this series; because two principal components are typically needed to explain at least 80% of the total phenotypic variation, an increased number of subsequent regression tests and adjustments for multiple comparisons are necessary. For individual level-data, among classical methods, MANOVA had a better statistical power compared to others, including GEE, LME, and OB (Table 5). With individual-level genotype and phenotype data, our proposed methods (dLC and eLC) demonstrated comparable power to MANOVA under both alternative hypotheses. However, dLC had less power than eLC in this scenario, likely due to the increased degree of freedom of its underlying chi-square distribution. Additionally, both of our methods outperform OB using aggregated summary test statistics when individual-level data are not available.

**Table 5.**
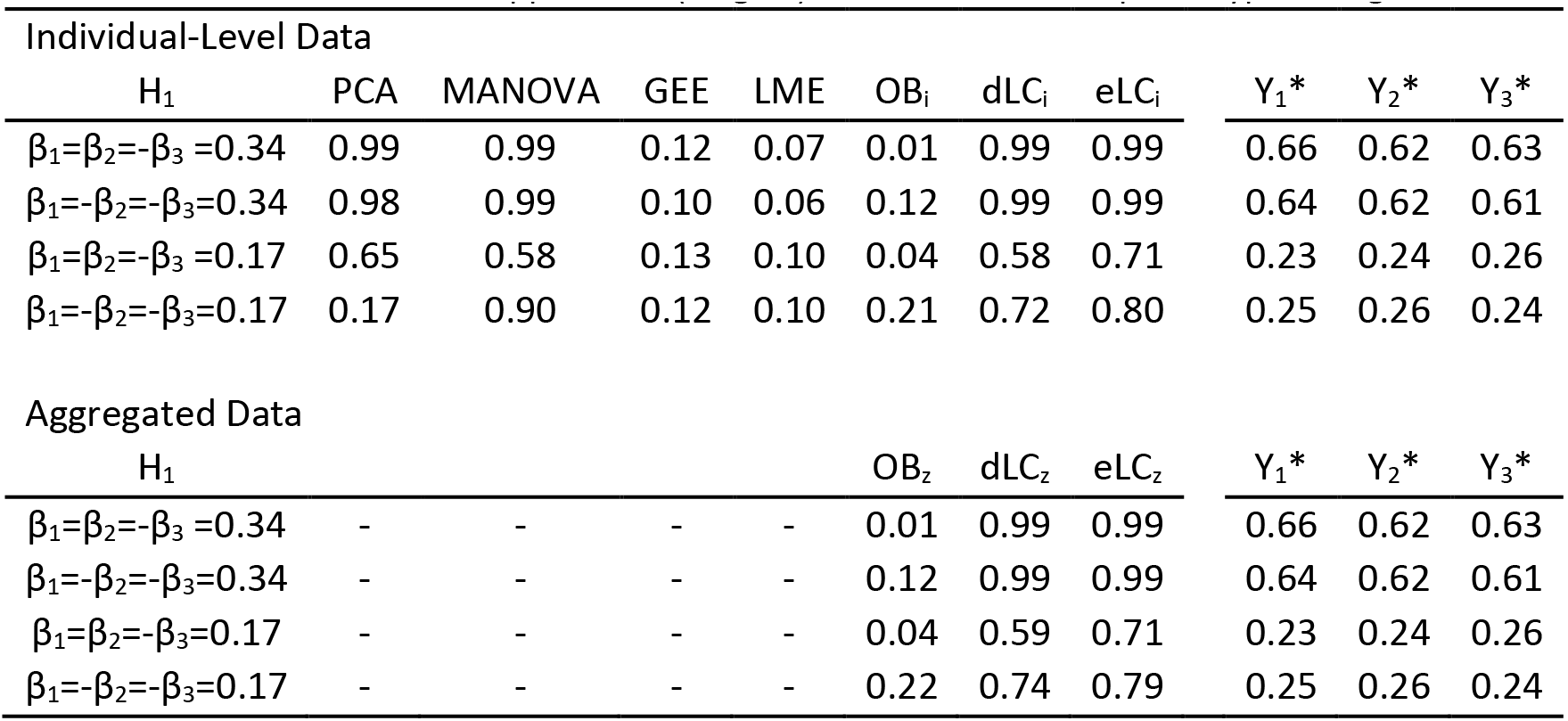
Power of multivariate approaches (Stage 1) on three correlated phenotypes using individual-level data and aggregated data

| Individual-Level Data |  |  |  |  |  |  |  |  |  |  |
| --- | --- | --- | --- | --- | --- | --- | --- | --- | --- | --- |
| H<sub>1</sub> | PCA | MANOVA | GEE | LME | OB<sub>i</sub> | dLC<sub>i</sub> | eLC<sub>i</sub> | Y<sub>1</sub>* | Y<sub>2</sub>* | Y<sub>3</sub>* |
| β<sub>1</sub>=β<sub>2</sub>=-β<sub>3</sub>=0.34 | 0.99 | 0.99 | 0.12 | 0.07 | 0.01 | 0.99 | 0.99 | 0.66 | 0.62 | 0.63 |
| β<sub>1</sub>=-β<sub>2</sub>=-β<sub>3</sub>=0.34 | 0.98 | 0.99 | 0.10 | 0.06 | 0.12 | 0.99 | 0.99 | 0.64 | 0.62 | 0.61 |
| β<sub>1</sub>=β<sub>2</sub>=-β<sub>3</sub>=0.17 | 0.65 | 0.58 | 0.13 | 0.10 | 0.04 | 0.58 | 0.71 | 0.23 | 0.24 | 0.26 |
| β<sub>1</sub>=-β<sub>2</sub>=-β<sub>3</sub>=0.17 | 0.17 | 0.90 | 0.12 | 0.10 | 0.21 | 0.72 | 0.80 | 0.25 | 0.26 | 0.24 |
| Aggregated Data |  |  |  |  |  |  |  |  |  |  |
| H<sub>1</sub> |  |  |  |  | OB<sub>z</sub> | dLC<sub>z</sub> | eLC<sub>z</sub> | Y<sub>1</sub>* | Y<sub>2</sub>* | Y<sub>3</sub>* |
| β<sub>1</sub>=β<sub>2</sub>=-β<sub>3</sub>=0.34 | - | - | - | - | 0.01 | 0.99 | 0.99 | 0.66 | 0.62 | 0.63 |
| β<sub>1</sub>=-β<sub>2</sub>=-β<sub>3</sub>=0.34 | - | - | - | - | 0.12 | 0.99 | 0.99 | 0.64 | 0.62 | 0.61 |
| β<sub>1</sub>=β<sub>2</sub>=-β<sub>3</sub>=0.17 | - | - | - | - | 0.04 | 0.59 | 0.71 | 0.23 | 0.24 | 0.26 |
| β<sub>1</sub>=-β<sub>2</sub>=-β<sub>3</sub>=0.17 | - | - | - | - | 0.22 | 0.74 | 0.79 | 0.25 | 0.26 | 0.24 |

*: p-values were adjusted using Bonferroni correction for these univariate analyses.

### Stage 2: Causal inference of the Pleiotropy

We utilized the Simulation Series II dataset (low effect size described in Table 1) to act as positive and negative controls in evaluating the performance of our proposed stage 2 method of testing pleiotropy under relevant hypotheses.

A specific null hypothesis, “β_1_>0,β_2_=β_1_+ε”, was simulated and used as the negative control. Table 6 presents the estimated type I error rates and power of our proposed method (the conditional testing of pleiotropy, cPLT) in comparison with the performance of the two classical conditional models strategy. Under the null hypothesis, cPLT demonstrated reasonable type I error rates, regardless of phenotypic residual correlations and effect directions.

**Table 6.** Type 1 error rates and power of the proposed Stage 2: causal inference of pleiotropy effects using individual-level data under relevant hypotheses

| $\rho^{\dagger}$ | Power<br>$\beta_1 = \vert \beta_2\vert > 0$ | | | Power<br>$\beta_1 > \vert \beta_2\vert > 0$ | | | Type 1 error<br>$\beta_1 > 0, \beta_2 = \beta_1 + \varepsilon$ | | |
| --- | --- | --- | --- | --- | --- | --- | --- | --- | --- |
| | cPLT | $Y_{c1}$ | $Y_{c2}$ | cPLT | $Y_{c1}$ | $Y_{c2}$ | cPLT | $Y_{c1}$ | $Y_{c2}$ |
| $\alpha = 0.05$ | | | | | | | | | |
| -0.50 | 0.150 | 0.263 | 0.259 | 0.199 | 0.536 | 0.050 | 0.065 | 0.364 | 0.050 |
| -0.25 | 0.495 | 0.441 | 0.413 | 0.306 | 0.566 | 0.089 | 0.049 | 0.349 | 0.066 |
| 0.01 | 0.789 | 0.608 | 0.595 | 0.487 | 0.618 | 0.179 | 0.066 | 0.344 | 0.047 |
| 0.25 | 0.522 | 0.417 | 0.389 | 0.314 | 0.538 | 0.086 | 0.053 | 0.349 | 0.055 |
| 0.50 | 0.151 | 0.260 | 0.270 | 0.183 | 0.504 | 0.050 | 0.045 | 0.326 | 0.046 |

Note: Y_c1_: conditional analysis for trait 1 adjusting for trait 2; Y_c2_: conditional analysis for trait 2 adjusting for trait 1.

^¶^: opposite effect direction was assigned when negative phenotypic correlation ρ was used.

Two alternative hypotheses, “β_1_= |β_2_|>0” and “β_1_> |β_2_|>0”, served as positive controls for detecting pleiotropy. cPLT demonstrated better performance than classical conditional strategies under these two alternative hypotheses (Table 6). However, the power was low for most methods when the degree of phenotypic correlations was high, suggesting the difficulty of using cross-sectional data to causally inference pleiotropy effects when phenotypes are highly correlated. Superior power was consistently found using cPLT under the β1= |β2|>0 alternative hypothesis, in which the genetic variant contributed to phenotypes equally and the phenotypic correlation was modest. In general, statistical power was highest when phenotypic residual correlation was very small, e.g., ρ =0.01. Further, power of cPLT decreased as the degree of phenotypic residual correlation increased. Interestingly, power did not seem to be as strongly influenced by the mixture of protective and deleterious genetic effects in this stage as it was in the GWAS screening stage (first stage).

These findings show that our proposed cPLT method performs well under the intended hypothesized pleiotropic relationship between genetic variants and correlated phenotypes, but its power can vary significantly as the correlation of phenotypes changes.

### Joint stage 1 and 2 reducing type 1 error of identifying pleiotropic effects

We further extensively examined the robustness of our proposed two-stage strategy under the null hypothesis, β_1_>0, β_2_=β_1_+ε (no pleiotropic effects). Specifically, we presented type 1 error rates for Stage 1 alone, Stage 2 alone and Stage 1 + Stage 2 with respect to various alpha levels (Table 7). Type 1 error rates were inflated for dLC applied alone in Stage 1, as expected. However, applying the proposed cPLT approach in Stage 2 would efficiently reduce the false positive results introduced in Stage 1. Notably, the overall false-positive rate was better controlled at the level of 0.05 when the screening threshold at Stage 1 was chosen at 0.025. Thus, this two-stage strategy would allow us to use **less-stringent cut-off p-values** in the Stage 1 GWAS screening to increase our power while effectively maintaining the overall false-positive rates through Stage 2.

**Table 7.** Type 1 error rates of separate and joint Stages under the null hypothesis, β_1_>0,β_2_=β_1_+ε

| $\alpha$ | 0.005 | | | 0.01 | | | 0.025 | | | 0.05 | | | 0.1 | | |
| --- | --- | --- | --- | --- | --- | --- | --- | --- | --- | --- | --- | --- | --- | --- | --- |
| $\rho^{\dagger}$ | S1 | S2 | Pool | S1 | S2 | Pool | S1 | S2 | Pool | S1 | S2 | Pool | S1 | S2 | Pool |
| -0.5 | 0.209 | 0.065 | 0.035 | 0.287 | 0.065 | 0.042 | 0.415 | 0.065 | 0.055 | 0.531 | 0.065 | 0.061 | 0.652 | 0.065 | 0.065 |
| -0.25 | 0.211 | 0.069 | 0.036 | 0.27 | 0.069 | 0.042 | 0.398 | 0.069 | 0.051 | 0.512 | 0.069 | 0.064 | 0.636 | 0.069 | 0.069 |
| 0.01 | 0.195 | 0.066 | 0.028 | 0.261 | 0.066 | 0.038 | 0.402 | 0.066 | 0.053 | 0.522 | 0.066 | 0.061 | 0.628 | 0.066 | 0.063 |
| 0.25 | 0.195 | 0.053 | 0.034 | 0.273 | 0.053 | 0.039 | 0.407 | 0.053 | 0.05 | 0.504 | 0.053 | 0.053 | 0.628 | 0.053 | 0.053 |
| 0.5 | 0.199 | 0.045 | 0.032 | 0.264 | 0.045 | 0.036 | 0.374 | 0.045 | 0.044 | 0.475 | 0.045 | 0.045 | 0.600 | 0.045 | 0.045 |

Note: S1: Stage 1 only; S2: Stage 2 only; Pool: pooled Stage 1+2; p-value=0.05 was fixed at Stage 2.

^¶^: opposite effect direction was assigned when negative phenotypic correlation ρ was used.

### Type 1 errors when filtering out SNPs with multivriate p-value lager than univariate p-values from each phenotype

Theoretically, the multivariate p-value of a true pleiotropic marker is considered to be smaller than that of its univariate results. We investigated this hypothesis using our simulation data under different alternative hypotheses. The proportions of SNPs with smaller p-values (an order of magnitude) in eLC under various scenarios are provided in Supplementary **Table S3**. In general, most p-values from eLC were smaller when the marker equally contributed to multiple traits and the phenotypic correlation was modest. On the other hand, p-values from eLC did not tend to be smaller when the phenotypic correlation was high or unequal genetic effect sizes existed. Adopting the smaller multivariate p-value filtering criteria added little improvement in terms of type 1 error. We further investigated the impact on statistical power by using this filter under relevant alternative hypotheses (Table 9). Unexpectedly, applying the smaller multivariate p-value filtering criteria would introduce modest to significant power loss, largely depending on the magnitudes of phenotypic correlations and genetic effect sizes.

**Table 9.** Power of multivariate approaches after filtering SNPs with multivariate p-values smaller than univariate p-values under relevant alternative hypotheses

|  | Simulation Series I. Higher Effect Size |  |  |  |  | Simulation Series II. Lower Effect Size |  |  |  |  |
| --- | --- | --- | --- | --- | --- | --- | --- | --- | --- | --- |
| | $\rho$ | dLC<sub>i</sub> | | dLC<sub>z</sub> | | $\rho$ | dLC<sub>i</sub> | | dLC<sub>z</sub> | |
|  |  | Full | Filter | Full | Filter |  | Full | Filter | Full | Filter |
| Equal Effect Size | -0.75 | 0.74 | 0.06 | 0.74 | 0.06 | -0.50 | 0.44 | 0.05 | 0.44 | 0.06 |
|  | -0.25 | 0.93 | 0.86 | 0.93 | 0.85 | -0.25 | 0.52 | 0.22 | 0.52 | 0.23 |
|  | 0.01 | 0.98 | 0.98 | 0.98 | 0.97 | 0.01 | 0.61 | 0.43 | 0.62 | 0.45 |
|  | 0.25 | 0.93 | 0.86 | 0.93 | 0.87 | 0.25 | 0.46 | 0.21 | 0.46 | 0.21 |
|  | 0.75 | 0.75 | 0.07 | 0.74 | 0.07 | 0.50 | 0.40 | 0.04 | 0.40 | 0.03 |
| Unequal Effect Size | -0.75 | 0.65 | 0.08 | 0.65 | 0.05 | -0.50 | 0.30 | 0.05 | 0.30 | 0.05 |
|  | -0.25 | 0.77 | 0.53 | 0.75 | 0.48 | -0.25 | 0.28 | 0.05 | 0.29 | 0.06 |
|  | 0.01 | 0.91 | 0.85 | 0.94 | 0.91 | 0.01 | 0.36 | 0.13 | 0.35 | 0.13 |
|  | 0.25 | 0.79 | 0.53 | 0.79 | 0.53 | 0.25 | 0.29 | 0.05 | 0.29 | 0.06 |
|  | 0.75 | 0.65 | 0.06 | 0.65 | 0.06 | 0.50 | 0.28 | 0.04 | 0.29 | 0.03 |

<sup>¶</sup>: opposite effect direction was assigned when negative phenotypic correlation $\rho$ was used.

*significance level was defined as p-value=1e-4 for higher effect size Simulation Series I and 0.01 for lower effect size Simulation Series II.

^¶^: opposite effect direction was assigned when negative phenotypic correlation p was used.

### An Application to GAW16 Framingham Heart Study data

To validate our newly developed two-stage approach to identify potential pleiotropic genetic effect, we applied our method to the GAW16 Framingham Heart Study dataset. We sought to identify pleiotropic variants (SNPs) associated with both serum high-density lipoprotein (HDL) and triglycerides (TG), known to influence cardiovascular disease. The phenotypic correlation for HDL and TG was -0.28, and the estimated correlation of univariate GWAS test statistics (~550K SNPs across genome) of HDL and TG was -0.32. A unified genome-wide significance level was defined by false discovery rate (FDR) at p-value=3*10^−6^, equivalent to q=0.05, for both bivariate and univariate analysis to screen potential pleiotropic SNPs. The genomic control inflation factor, 1GC was 1.01 for this bivariate GWAS analysis. Q-Q plot of the bivariate GWAS results (**Supplementary Figure S1**) showed no inflation beyond that expected by chance alone. A further comparison of bivariate results from eLC between individual-level data and aggregated summary statistics (**Supplementary Figure S2**) revealed no substantial differences.

We identified 25 genome-wide significant bivariate loci from the stage-1 GWAS screening using aggregative statistics. We then selected the most significant SNP in each locus to perform our stage 2 cPLT-analysis using individual level data for causal inference of pleiotropic effects. The cPLT method with 10,000 permutations were performed (Table 8). The significance level in stage 2 was defined as p < 0.002 after Bonferroni correction. Our two-stage analysis identified 2 SNPs, rs3200218 in the coding region of *LPL* gene and rs8192719 on the exon/intron boundary of *CYP2B6* gene. These two loci served as positive controls in the GAW16 simulation dataset, and were validated as pleiotropic genes using our strategy. The simulated heritability of rs3200218 for HDL and TG was 0.3% and 0.4%, respectively. For rs8192719, the heritability for HDL and TG was reported as 0.3% for both [42]. Additionally, as a negative control, SNP rs7031748 demonstrated a significant p-value from bivariate GWAS analysis in the Stage-1 screening and it failed to reject the null hypothesis in the Stage-2 cPLT analysis (causal inference), suggesting some of the significant bivariate associations of genetic variants in the stage-1 GWAS screening likely resulted from indirect correlations among phenotypes due to non-genetic effects, rather than a causal pleiotropic relationship; therefore, the stage 2 is essential to identify pleiotropic genetic effects.

**Table 8.** Selected pleiotropic SNPs associated with HDL and TG in GAW16 simulation dataset after applying proposed two-stage approach

| rsSNP | MAF % | Gene | Univariate |  | dLC P value | Conditional |  | cPLT P value |
| --- | --- | --- | --- | --- | --- | --- | --- | --- |
|  |  |  | HDL | TG |  | HDL | TG |  |
| rs3200218 | 21.7 | LPL | 8.08E-08 | 4.83E-04 | 1.34E-10 | 1.28E-11 | 5.74E-08 | <1E04 |
| rs8192719 | 24.9 | CYP2B6 | 7.08E-06 | 8.17E-06 | 2.94E-06 | 9.91E-04 | 7.02E-04 | <1E04 |
| rs7031748 | 12.7 | ABCA1 | 5.59E-09 | 0.725 | 1.76E-07 | 4.24E-10 | 0.025 | 0.014 |

## Discussion

In this study, we propose a novel two-stage approach to identify potential pleiotropic effects on a genome-wide scale. Using powerful linear-combination approaches as a screening tool, we demonstrated the plausibility of effectively utilizing summary statistics (without individual level of genotypes and phenotypes), e.g., univariate GWAS meta-analysis results, to perform multivariate analysis and screen potential pleiotropic effects in the first stage. Comparing to classical multivariate methods and previously proposed methods by others, the proposed methods (eLC and dLC) consistently demonstrate better power, maintain expected type 1 error rates, and do not require complex modeling assumptions in most of the circumstances. More important, the proposed methods show considerable power over other approaches when mixed genetic effects and opposite effect directions are existing in the tested phenotypes. Based on simulations, our proposed causal inference approach for the pleiotropic effects (cPLT method) in the stage-2 provided reasonable power while maintaining conservative type 1 error rates when traits were moderately correlated. We are able to leverage univariate GWAS summary statistics and improve power to identify pleiotropic effects using our two-stage design. We are able to validate our approach by identifying the positive controls of pleiotropic effects in the GAW16 simulation datasets [42].

Meta-analyses that synthesize results from different studies have become a standard approach to increase statistical power in many GWAS. Most multivariate methods, however, require individual level of data and cannot utilize summary statistics of GWAS meta-analysis that most of them are publicly available. Our novel strategy can aid the discovery of pleiotropy in meta-analysis of GWAS results. While traditional multivariate approaches, such as PCA and MANOVA, are powerful when individual-level data are accessible, our simulation studies demonstrate the effectiveness of our proposed eLC and dLC approaches when summary test statistics are available. The use of summary test statistics is advantageous for reducing potential bias introduced by population admixture or stratification, given that these confounders have been well adjusted in genome-wide association analyses in each participating study. In addition, instead of using the covariance matrix estimated through individual-level data as O’Brien suggested, we demonstrate the feasibility of using the covariance of summary test statistics to effectively increase the power as in a meta-analysis. The proposed methods can be applied to population-based studies and familial data; as well as quantitative traits and binary traits. For family-based studies, we also provided a simple algorithm (see Supplementary material for detailed description) for estimating covariance matrix in individual-level data.

Many factors affect statistical power of the proposed approach. The statistical power of genetic analyses relies on the underlying genetic effect sizes. To evaluate the impact of genetic effect size, we specifically simulated SNPs with smaller effect sizes (Simulation Series II). Our proposed methods were able to detect the pleiotropic effects. Substantial power can also be gained by using the proposed dLC and eLC approaches when the genetic effects are not in the same direction. Interestingly, eLC shows superior power to dLC when more than two traits are combined, despite similar performance to dLC in bivariate scenarios. Considering the intensive computational process involved in applying eLC, for bivariate GWAS analysis (only involving two phenotypes), dLC provides us a less computational intensity way to mine potential pleiotropic effects. For multivariate GWAS analysis with more than five phenotypes involved, eLC is preferable. Another important factor determining power is minor allele frequency (MAF) of the genetic marker of interest. We fixed MAF at 10% in all our simulation studies. Notably, however, the relative efficiency of multivariate methods seems to remain consistent but the degree of power changes as MAF varies [26,37]. The results presented here should be comparable when the proposed methods are applied to genetic variants with other MAFs.

Several multivariate methods (required individual level of data) have been proposed by others, such as the canonical correlations analysis (CCA) [43] implemented in PLINK[44]. Our simulation showed that this approach has similar power, as expected, to the MANOVA in all the scenarios (data not shown). Multivariate approaches for family data, such as the FBAT-GEE and FBAT-PC, proposed by Lange et al. [20,45] were also evaluated. Our approach outperformed these methods (data not shown). FBAT-based approaches usually introduce concerns about a substantial loss in power, since between-family correlations are ignored in the nature of an FBAT algorithm. In our simulations, FBAT has far less power to detect main effect markers compared to the LME model, the loss of power may be exaggerated in the context of combining summary statistics on a genome-wide scale, making it difficult to achieve a significance level that supports a positive finding for multivariate analysis. Different from our proposed approaches, eLC and dLC, above methods cannot be used when only aggregative summary statistics are available, suggesting a broader application that our approaches can be applied.

The proposed method of testing pleiotropy at the stage 2, cPLT method, allows us to test the specific alternative hypothesis that a genetic marker of interest is independently associated with multiple phenotypes, rather than the typical alternative hypothesis that association with one phenotype is sufficient in multivariate methods. In our simulations, we significantly improve the power to infer true pleiotropy over the two classical conditional modeling strategies [40]. However, there are limitations of our proposed cPLT method. The proposed cPLT method of testing pleiotropy appears to be conservative when phenotypes are closely correlated. In an extreme case, when two traits are nearly unrelated, meaning γ_1_ ≈ 0, the individual score in equation (5) (see Methods) will be very similar to the regular regression between X and Y. On the other hand, two traits are statistically reduced to one when they are closely correlated. Hence, there is little independent variance left in the proposed individual score in equation (5). This is a common problem for all causal inference statistical algorithms when dealing with highly-correlated outcomes or phenotypes. Therefore, as for many association analyses, additional biological evidence or animal studies are needed to validate our statistical detection of pleiotropy.

It is important to note that the selection of cut-off value in the Stage-1, multivariate GWAS screening stage, influences the overall false-positive findings in the two-stage approach. It will be too conservative when a stringent threshold (such as the commonly used genome-wide significant threshold, p < 5x10^−8^) is applied. We suggest apply a commonly used genome-wide suggestive threshold, such as p < 5x10^−6^, however, further studies are warranted to investigate the best choice of threshold in the multivariate GWAS screening stage (Stage-1) to maximize overall power and maintain reliable type 1 error. In addition, adopting the smaller multivariate p-value filtering criteria (smaller multivariate p-values comparing to univariate p-values) added little improvement in terms of type 1 error, but had a significantly reduced power. The power loss was depended on the magnitudes of phenotypic correlations and genetic effect sizes.

In summary, we have developed a novel and powerful two-stage approach to identify pleiotropic effects on a genome-wide scale without complex modeling assumptions. Our extensive simulations consistently demonstrated the advantages of the proposed approach over other existing multivariate analyses or methods to identify pleiotropic effects. The stage-1 multivariate GWAS screening in our two-stage approach does not need to have individual level of genotypes and phenotypes, which extend a broader application. By leveraging GWAS meta-analysis results, we can efficiently and effectively screen for potential pleiotropic markers on a genome-wide scale. A candidate marker set then can be selected for the second stage to test of independence, thus confirming the “real” pleiotropic effects. A computational program written in C++ for our proposed combining approach (eLX) is available at https://sites.google.com/site/multivariateyihsianghsu/.

## Methods

We proposed to identify pleiotropic effects on a genome-wide scale through a two-stage design (Figure 1). Briefly, at stage 1, we developed two powerful combining test statistics methods to screen for potential pleiotropic-effect SNPs on a genome-wide scale. At stage 2, a candidate marker set was then derived from Stage 1 and tested using a novel approach of testing pleiotropy, called conditional pleiotropy testing (cPLT).

### Stage 1

#### Direct linear combination of correlated test statistics (dLC)

In general, let **T**= (T_1_, T_2_,…,T_k_)^T^ denote a vector of K correlated test statistics obtained individually from each univariate analysis for a specific trait against a genetic marker. Under most circumstances in current GWAS studies, T usually follows an asymptotical multivariate normal distribution with mean **β** = (*β_1_, β_2_, … β_k_*)^T^ and known or estimated covariance **Σ**, where **Σ** is a k × k symmetric matrix. Assume the null hypothesis we want to test in multivariate analysis is H_0_: **β** = 0. In other words, the genetic variant is not associated with any phenotype. In contrast, the general alternative hypothesis H_1_ is at least one *β_k_* >0, k=1,…,k. Extending from O’Brien’s theory[33], we propose a new approach for combining correlated traits, called direct linear combination of test statistics (dLC). The new test statistics of dLC can be written as:

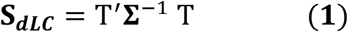

Under the null hypothesis, **S**_*dLC*_ follows a χ^2^ distribution with K degrees of freedom and can be effectively used to test the joint significance of dependent univariate test statistics.

#### Empirical linear combination of correlated test statistics (eLC)

As illustrated in Xu et al. [33], dLC may not have optimal power against specific alternatives resulting from the heavy tail of the χ^2^ distribution. Therefore, we further proposed a data-driven empirical approach to combine correlated test statistics as:

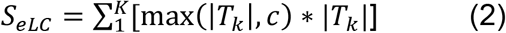

where c is some given non-negative constant. The weight in this new test statistics will be optimally determined by the specific data structure. For instance, when c=0, the test statistics simply reduces into sum of squares of T_k_. When c is relatively large, equal weight is assigned to each T_k_. Ideally, we would like to find an optimal value of c, so the *S_eLC_* performs as a linear combination of T_i_ when under the H_0_; but, under the alternative H_A_, more weight is given to the larger true T_i_. The bona fide p-value for *S_eLC_* then can be estimated by applying permutation or perturbation techniques (see Supplementary).

##### Estimation of the Variance-Covariance Matrix **Σ**

Several methods can be used to estimate the covariance matrix **Σ** of univariate test statistics. For simplicity, we demonstrate two estimation approaches in a bivariate scenario. Based on the method used in Yang et al. [37], we first utilize the sample covariance matrix of the test statistics of all SNPs from univariate GWAS analyses as an approximation. **Σ**, the covariance matrix, thus can be estimated as:

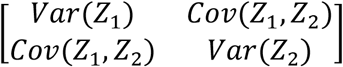

where Z_1_ consists of unbiased univariate test statistics of all the SNPs for the first trait on genome-wide scale, so does Z_2_. On the other hand, **Σ** can be estimated by using generalized least squares from individual-level data, as suggested by O’Brien [34]. A similar approach is also demonstrated by Liu et al. [26].

In this paper, the combining tests utilizing a covariance matrix approximated from the summary GWAS test statistics will be denoted by OB_z_, dLC_z_, and eLC_z_. Similarly, those with a covariance matrix calculated from individual-level data will be referred to as OB_i_, dLC_i_, and eLC_i_.

##### Simulation Study

Monte-Carlo Simulations were employed to generate data for evaluating the validity and performance of all the multivariate methods, especially the proposed dLC and eLC approaches. The main scenarios and key parameters for simulating genotype data are shown in Table 1. Various conditions were considered, including equal genetic marker contributes to both traits (Scenario A) and unequal genetic effects (Scenario B). To evaluate the proposed pleiotropy test strategy in Stage 2, a special Scenario C was also introduced. In particular, a single continuous trait was first generated with an assigned effect size. The second trait was then simulated by artificially adding random noise on the first generated trait. Therefore, the genetic variant is directly associated with the first trait but indirectly linked with the second trait, as illustrated in R2 and R3 in Figure 2. In all simulations, only quantitative traits and unrelated subjects were generated. A sample size of 1000 subjects was simulated in each of 1000 replicates.

**Figure 2.**
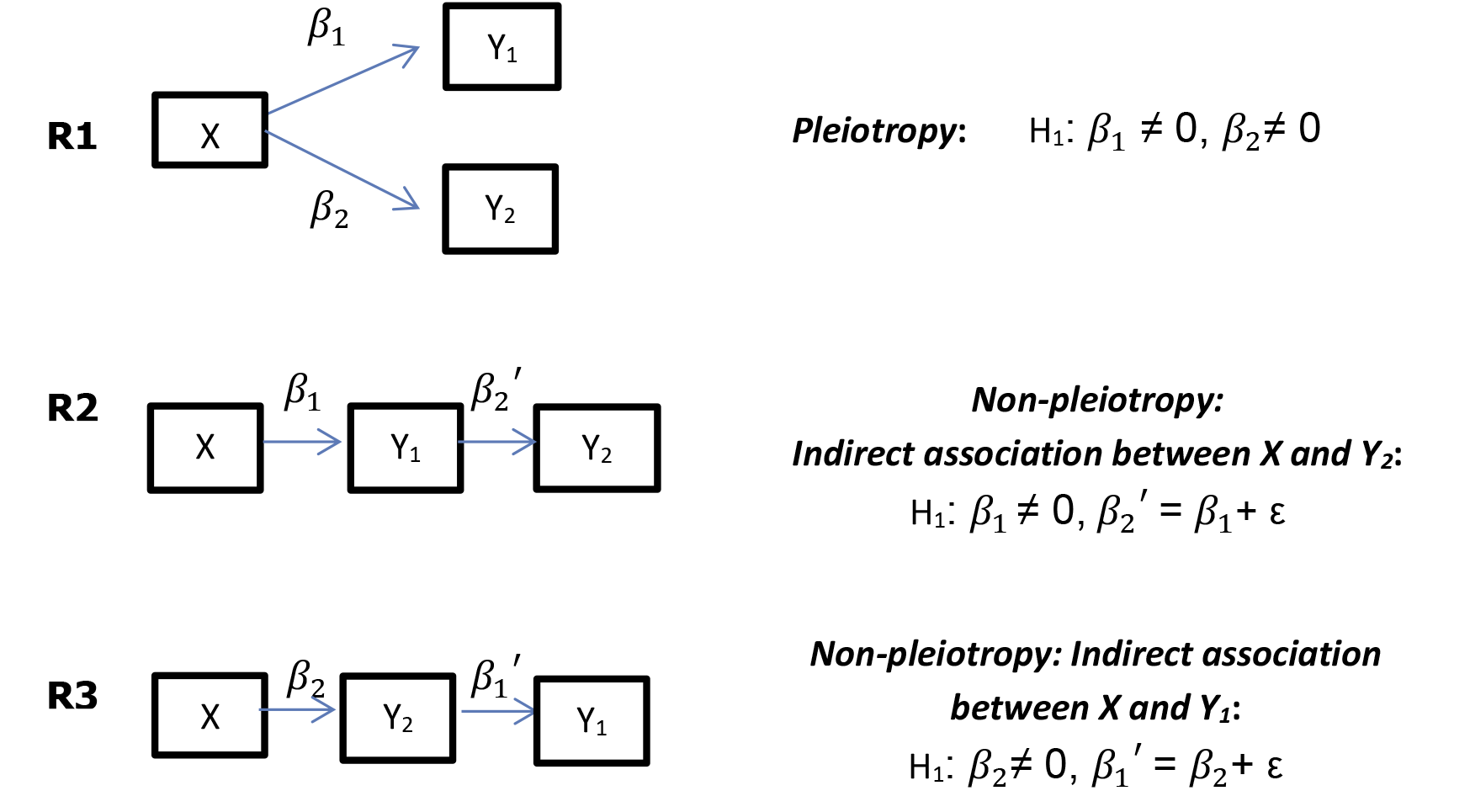
Possible relationships between genetic marker X and two correlated phenotypes Y_1_ and Y_2_ when the null hypothesis of a multivariate model is rejected. *β*_1_ represents direct genetic effects to phenotype Y_1_ contributed by X; *β_2_* represents genetic effects to phenotype Y_2_ directly contributed by X. *β*_1_ ’ and *β*_2_ ’ represent indirect associations between X and Y_1_ and Y_2_, respectively.

The effect size of the SNP on each trait is estimated from 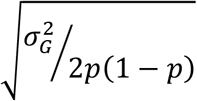, where p is the minor allele frequency (MAF) at the locus and 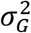, heritability, is the phenotypic variance explained by the SNP. The heritability is simulated at 1% and 2% in the Simulation Series I, and 0.5% and 0.1% in Simulation Series II for each trait, respectively. The variance of the environmental effects was fixed as 1 in all simulation studies. The genotypes were then generated under Hardy-Weinberg equilibrium with a specified MAF at 10%.

Bivariate quantitative phenotypes were randomly drawn from a bivariate normal distribution to represent the pleiotropic relationship R1 in Figure 1:

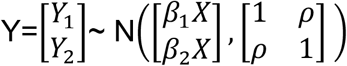

where *β_i_* is the additive genetic effect size for trait i, X is the additive score of the coded allele, and *ρ* is the residual correlation between Y_1_ and Y_2_, respectively.

Various simulation scenarios with respect to *ρ* were further generated. Briefly, *ρ* was selected at -0.75, -0.25, 0.01, 0.25, and 0.75 to mimic the correlations we have observed in real GWAS data analyses in the Simulation Series I. A smaller range of *ρ* at -0.5, -0.25, 0.01, 0.25 and 0.5 was also employed in Simulation Series II with lower genetic heritability. Notably, a mixture of protective and deleterious genetic effects was introduced when negative *ρ* was used in generating data. For instance, we assigned β_1_=- β_2_ as the alternative hypothesis when *ρ* was -0.25.

In addition to bivariate scenarios, we also generated three moderately correlated quantitative traits to evaluate the performance of various approaches when more than two traits were combined. For simplicity, the genetic heritability was equivalently set at 0.5% for each of the three traits. And the pairwise phenotypic residual correlation was chosen at 0.25, -0.15, and -0.20. The data were then generated under the null hypothesis H_0_: β_1_=β_2_=β_3_=0 and two alternative hypotheses, H1: β_1_=-β_2_=-β_3_ >0 and H_1_: β_1_=β_2_=-β_3_ >0, respectively.

Each replicate was analyzed by using OB, dLC, and eLC approaches individually. We estimated the covariance matrix **Σ** through approximation from summary data and individual-level data. Other multivariate methods were also compared, including LME with a random effect accounting for phenotypic correlations, GEE, MANOVA, and PCA, in which the first component was used as a dependent variable in the subsequent linear model. All analyses were conducted in R software (http://r-project.org/). Power and type I error rates of each approach were calculated as the proportion of replicates with a p-value less than a given significant threshold in the corresponding scenarios. Specifically, power was derived with 1000 replicates with the significance level at p-value equal to 10^−4^ for each true scenario in Simulation Series I, and 10^−2^ for those in Simulation Series II. Type I error rates were estimated in the settings that β_1_=β_2_=0 with 1000 replicates at nominal significance levels of 0.05 and 0.01. In the context of adjusting multiple testing for two univariate association tests of two phenotypes, standard Bonferroni corrections were applied with the significance level at 5*10^−5^ and 5*10^−3^ for Simulation Series I and II separately.

##### Stage 2, Testing of Pleiotropy

The null hypothesis of multivariate approaches, H_0_: β_k_=0 (j=1, 2.,.k), is rejected when at least one of β_i_ is not equal to 0. This would result in three possible alternative relationships for bivariate scenarios (Figure 1). Briefly, when two traits are linked and only one of them is indeed associated with the genetic marker, it is likely the other trait will be indirectly associated with the same marker even in absence of real pleiotropy, as demonstrated in R2 and R3. Pleiotropy explained in R1, however, will be true only when the genetic marker is independently associated with two traits under the alternative hypothesis, H_1_:β_1_≠0 and β_2_≠0.

Let Y_i_ and K_i_ be the target phenotypes for the *i*th subject in the study and X_*i*_ be the genetic marker of interest for this subject. A regular strategy of testing this specific hypothesis involves applying two separate conditional models for each trait with adjusting the other one as a covariate. Specifically, the following linear model is usually used to test the direct genetic effect of marker X contributing to quantitative phenotype Y_i_

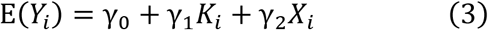

where γ_0_,γ_1_, and γ_2_ denote the mean parameters can be estimated by ordinary least squares approach.

Here, we proposed a new approach of testing pleiotropy by combining adjusted phenotypes together based on the framework suggested in Lange et al. [40].

From equation (3), we can adjust the phenotype Y_*i*_ for the effects contributed by the phenotype K_*i*_ for *i*th subject as:

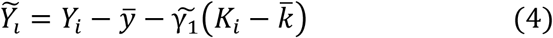

where 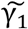 is the ordinary least squares estimate for γ_l_, and 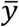 and 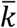 represent the observed sample mean of phenotype Y and K, respectively. In a population-based study, we can derive a new test statistic for 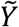 contributed by all subjects as:

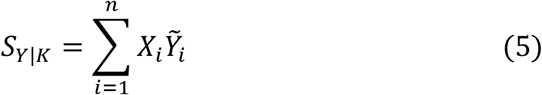

Likewise, we can construct the new test statistics for 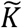 as: 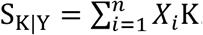. Both S_Y|K_ and S_Y|K_ can be considered as direct scores between the genetic marker and the target phenotype. To test the specific alternative hypothesis of pleiotropy, we further proposed a new approach as

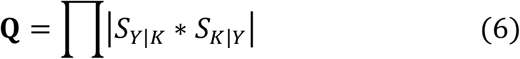

This new test statistics Q can only be rejected when both adjusted individual statistics are significant (β_K,X|Y_≠0 and β_Y,X|K_≠0). The empirical p-value of Q can be estimated by applying a permutation test.

We have conducted comprehensive studies to evaluate the performance of this novel approach of inferring pleiotropy in the Simulation Series II dataset. Power was defined as the number of significant results with p-values less than 0.05 and 0.01 in 1000 replicates. The robustness of this approach and two-stage strategy were further examined under a specific null hypotheses, β_1_>0,β_2_=β_1_+ε. The performance of the two conditional models strategy was also investigated and compared to our proposed method.

##### GAW16 Problem 3 simulation data sets

The first replicate in the Genetic Analysis Workshop 16 (GAW 16) Problem 3 simulation data sets [42] was selected to validate our approach [42]. The GAW 16 derives from the Framingham Heart Study (FHS), which includes 6,476 subjects from 3 generations with real genotypes of approximately 550,000 SNP markers and six simulated phenotypes with multiple measurements. Subjects are distributed among 942 families and 188 singletons. The two directly-simulated quantitative traits, high-density lipoprotein (HDL) and triglycerides (TG), were used in our analysis. Genotypes were derived from the Affymetrix GeneChip Human Mapping 500K Array set and the 50K Human Gene Focus Panel. Standard analytical approaches for GWAS are illustrated in detail in the Supplementary.

## Annotations

***PCA:*** Principle Component Analysis; ***MANOVA:*** Multivariate Analysis Of Variance; ***GEE:*** Generalized Estimation Equations; ***LME:*** Linear Mixed Effects models; ***OB:*** O’Brien combining statistics method; ***dLC:*** Direct Linear Combining statistics method; ***eLC:*** Empirical Linear Combining statistics method; ***cPLT:*** conditional Pleiotropy Testing method

***S_dLc_:*** Test statistics for direct Linear Combining method

**SeLc:** Test statistics for empirical Linear Combining method

**HDL:** High-density lipoprotein; TG: Triglycerides

